# Vesalius: high-resolution in silico anatomization of Spatial Transcriptomic data using Image Analysis

**DOI:** 10.1101/2021.08.13.456235

**Authors:** Patrick C.N. Martin, Hyobin Kim, Cecilia Lövkvist, Byung-Woo Hong, Kyoung Jae Won

## Abstract

Characterization of tissue architecture promises to deliver insights into development, cell communication and disease. *In silico* spatial domain retrieval methods have been developed for spatial transcriptomics (ST) data assuming transcriptional similarity of neighboring barcodes. However, domain retrieval approaches with this assumption cannot work in complex tissues composed of multiple cell types. This task becomes especially challenging in cellular resolution ST methods. We developed Vesalius to decipher tissue anatomy from ST data by applying image processing technology. Vesalius uniquely detected territories composed of multiple cell types and successfully recovered tissue structures in high-resolution ST data including in mouse brain, embryo, liver, and colon. Utilizing this tissue architecture, Vesalius identified tissue morphology specific gene expression and regional specific gene expression changes for astrocytes, interneuron, oligodendrocytes, and entorhinal cells in the mouse brain.

## Main Text

From the smallest cell to the largest organ, we find patterns of organization supporting homeostasis within organisms. Each cell functions in the context of its neighbors and each group of structured cells functions in the context of tissue architecture. The investigation of this multi-level organization promises to deliver insight into development, cellular communication and disease.

One approach to probe into the organization of tissues and cells is by using Spatial Transcriptomics (ST)^1,2^. ST is a set of methods that recover gene expression while maintaining the spatial component intact. A number of ST methods have been developed in the last few years and fall into two categories: image-based approaches using fluorescence *in situ* hybridization (FISH) and sequencing-based approaches using spatially resolved barcodes. Image-based ST approaches including seqFISH^3,4^ or merFISH^5^ provide sub-cellular resolution ST. Image-based techniques rely on the pre-selection of target mRNA species and - due to the challenge of distinguishing overlapping fluorescent signals - are limited to a smaller number of genes sampled at a time^6^.

In contrast to image-based ST, sequencing based-ST techniques such as 10X Visium^7^, Seq-Scope^8^ or Slide-seq^9,10^ provide a non-biased and genome wide quantification of mRNA species. Technological advances have made it possible to obtain sequencing based-ST data on cellular and even sub-cellular resolutions. Tissue territory detection in high-resolution ST data will strengthen our understanding of tissue architecture and its associated marker genes providing further opportunities to study transcriptomic changes due to local environment and tissue morphology.

A number of tools including Seurat^11^, BayesSpace^12^ and Giotto^13^ have been developed to understand tissue architecture from ST data. Seurat leverages refence single cell data sets to map barcode identities to their respective location in ST data. While this approach demonstrates the cellular heterogeneity of tissues, the task of recovering and extracting anatomical regions is still challenging due to their cellular complexity. On the other hand, BayesSpace and Giotto provide distinct models both utilizing Hidden Markov Random Fields. Their respective methods attempt to cluster barcodes together under the assumption that neighboring barcodes are likely part of the same cluster if transcriptionally similar. These methods have performed well in low resolution Visium 10X data^12,13^. However, this assumption will not necessarily hold in complex tissue containing multiple cell types as neighboring barcodes are just as likely to represent a different cell type in high resolution ST data. The recovery of anatomical structures containing multiple cell types becomes extremely arduous and generally has relied on manual isolation. One solution to this challenge is to link anatomical territories from companion hematoxylin and eosin (H&E) staining images to their spatial transcriptomic assay^14–18^. However, techniques such as Slide-Seq^9,10^ do not provide these companion images. Furthermore, the segmentation of anatomical territories in images still remains challenging without manual annotation^19–21^.

To address these limitations, we developed Vesalius – an R package – designed to perform *in silico* anatomization and isolation of tissue territories without the use of H&E companion images. Vesalius converts the transcriptome into an RGB color code that is then embedded into an image. By leveraging a variety of image analysis techniques, Vesalius is able to recover complex tissue territories in the mouse brain, mouse embryo, diseased liver and colon. Comprehensive tests using simulated data indicate that Vesalius can identify tissue territories even when the tissue is composed of multiple cell types while other competitors developed for low-resolution ST data often identify numerous unwanted patches. Identified territories are used to discover tissue morphology-specific gene expression as well as changes in the transcriptome of astrocytes, interneuron, oligodendrocytes, and entorhinal cells in the mouse brain depending on the tissue territory.

## Results

### Vesalius embeds the transcriptome in the RGB color space

The core concept of the Vesalius algorithm is to represent the transcriptome of a barcode as a color in the RGB color space and build images upon which image analysis techniques can be applied (Fig.1a-b). To embed the transcriptome into images, Vesalius first pre-processes sequencing based spatial transcriptomic data by log normalizing and scaling counts values, extracting highly variable features, and reduces dimensionality via principal component analysis (PCA). Next, Vesalius uses Uniform Manifold Approximation and Projection (UMAP) to project PCs into three dimensions and embed the latent space into the RGB color space (See Methods – Fig. S1a). Alternatively, Vesalius can also embed PC loading values into the RGB color space for a more targeted view of data variance (See methods – Fig. S2). Vesalius handles the uneven location of barcodes in the ST assay by expanding punctual coordinates into multi-pixel tiles using Voronoi Tessellation. Images are constructed by associating color codes to their respective tile.

**Fig. 1.**
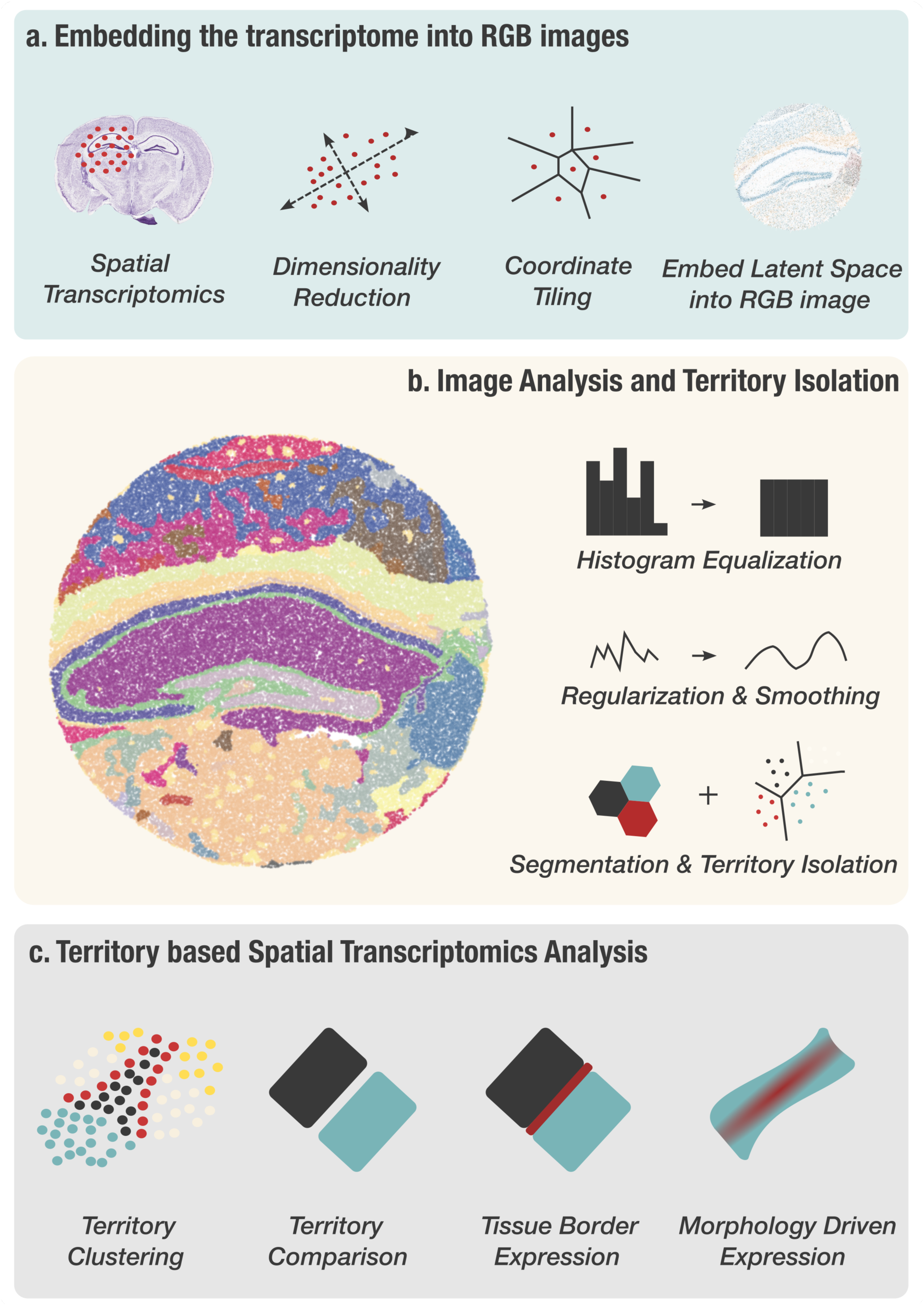
Describing Tissue anatomy with Vesalius. (**a**) Vesalius embeds ST data into RGB colored images. This is achieved by preprocessing ST data and reducing dimensionality. In parallel punctual ST coordinates are converted into tiles. Finally, the UMAP latent space (or PCA loading values) is transformed into an RGB color space and the color code attributed to each barcode is assigned to its respective tile. (**b**) Vesalius applies image analysis techniques to RGB images describing the transcriptional landscape of a tissue with the aim of isolating tissue territories. (**c**) Vesalius enables a territory-based ST framework including spatial territory clustering, territory comparison, tissue border expression, and morphology driven expression.

Next, Vesalius applies image analysis techniques in order to perform *in silico* anatomization (Fig. 1b). After balancing the color histogram and smoothing (see Methods), image segmentation based on k-means clustering is applied to produce color clusters that can be further subdivided into territories. Image processing parameters can be finely tuned depending on the user’s interest and the ST method used. Vesalius checks if barcodes are within a certain capture radius of each other. Vesalius will assign barcodes belonging to the same color cluster into separate territories if they are far enough from each other in 2D space (See Methods - Fig. S1b).

The isolation of anatomical territories and image representation of ST data with Vesalius enhances ST analysis by providing a territory-based framework (Fig. 1c). Isolated territories can be further clustered to recover the finer details of cellular organization. Territories can be compared to investigate territory specific gene expression. Neighboring territories can be manipulated to recover tissue border gene expression and gene expression patterns arising within specific anatomical structures.

### Vesalius overcomes the challenge of isolating anatomical structures containing heterogenous cell populations

High-resolution sequencing-based ST methods promise an unbiased and spatially resolved view of the transcriptome. Yet, most tissues contain multiple cell types and it can be challenging to recover uniform anatomical structures especially when no companion H&E images are provided. To demonstrate Vesalius’s ability to isolate anatomical structures, we used Slide-seq V2 that provides a high-resolution sequencing-based ST assay for mouse hippocampus and embryo^10^.

Vesalius identified 41 territories in the mouse hippocampus (Puck_200115_08) including the CA field, the Dentate Gyrus and the Corpus Callosum (Fig. 2a - Image is from Allen Institute^22^). Territories characterized by too little barcodes (<50) and too far away from another territory were described as being isolated. Territories recovered by Vesalius are generally characterized by uniform regions that match well with reference annotation (Fig 2a). In contrast to Vesalius, BayesSpace^12^ and Seurat^11^ recover anatomical structures insofar as these structures are characterized by a homogenous population of cells but produce numerous small patches potentially due to the heterogenous nature of the brain tissues (Fig. 2b, Fig. S3 and Fig. S4).

**Fig. 2.**
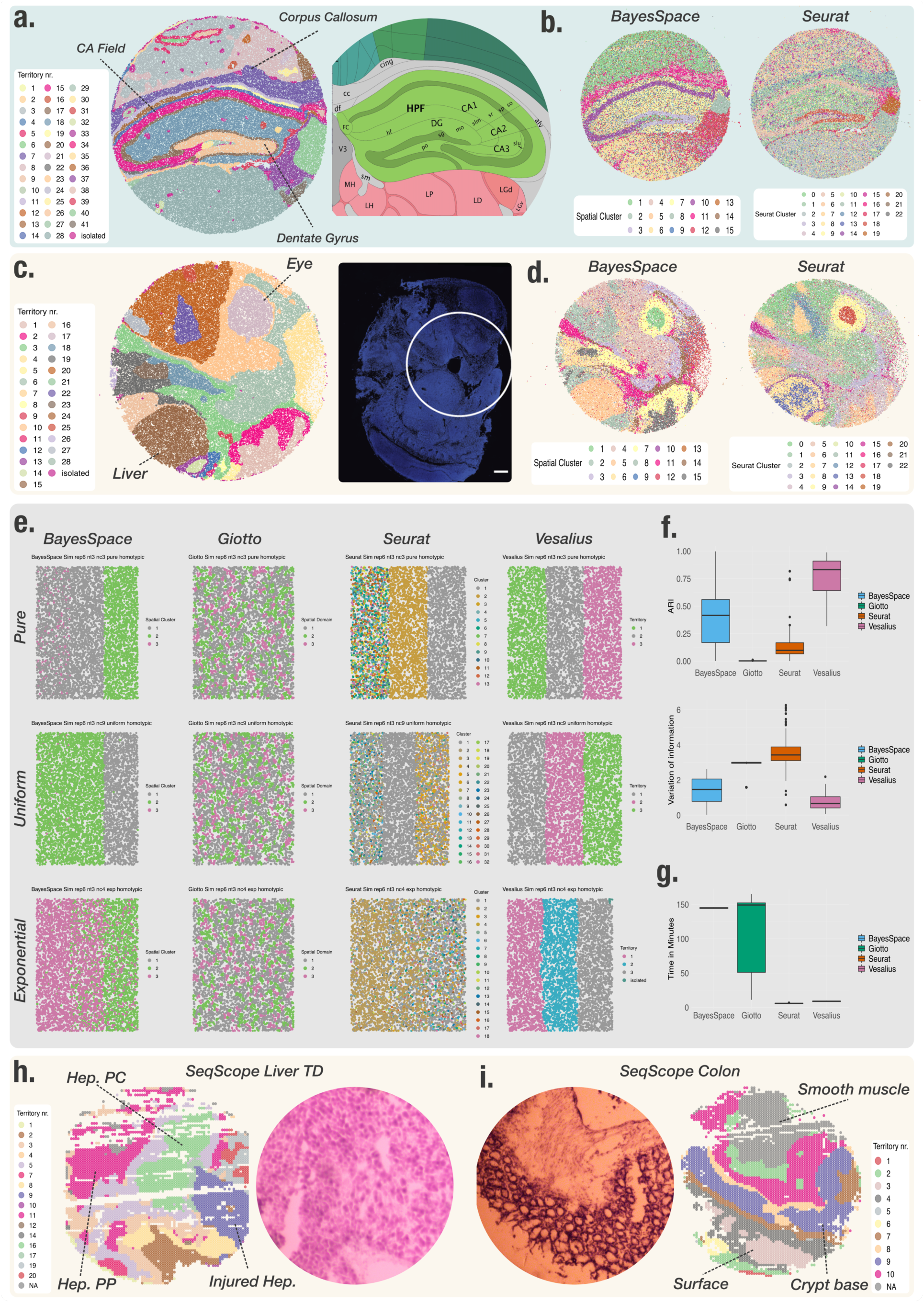
Vesalius recovers uniform anatomical territories in high resolution ST data. **(a)** Vesalius accurately recovers tissue territories in Slide-seqV2 data taken from the mouse hippocampus and surrounding brain (Puck_200115_08). A comparison with the Allen Brain Atlas reference atlas illustrates that Vesalius recovers many structures such as the Dentate Gyrus, Corpus Callosum and the CA field. **(b)** BayesSpace and Seurat applied to the same data set recover structures insofar as these structures contain homogenous cell populations. The identified clusters are dispersed over the entire tissue section and thus do not represent a clear tissue territory. **(c)** Vesalius recovered uniform tissue territories in the mouse embryo (Slide-seqV2 - Puck_190926_03). The microscopy image highlights the section of the embryo used to produce Puck_190926_03 (Image taken from Slide-seq V2^10^). **(d)** BayesSpace and Seurat failed in selecting specific tissue territories. (**e**) Example of simulated spatial territories retrieved by each tool under the pure regime (top row), the uniform regime (middle row – n = 3) and the exponential regime (bottom row – n = 4) in simulation replicate 6. (**f**) Vesalius outperforms BayesSpaces, Giotto and Seurat in simulated high-resolution ST. Top panel shows ARI scores and bottom panel shows Variation of Information. (**g**) Comparison of running time between each tool in simulated ST data. **(h)** Vesalius identified tissue territory using SeqScope early-onset liver failure (Sample 2117) and **(f)** healthy colon (Sample 2111). Vesalius highlights various hepatocyte populations (**e**) such as Pericentral Hepatocytes (Hep. PC), Periportal Hepatocytes (Hep. PP) and injured Hepatocytes (Injured Hep.). Territory isolation in the colon (i) shows various structures and layers including the smooth muscle, the crypt surface, and the crypt base.

We observed a similar phenomenon when we analyzed the mouse embryo Slide-seqV2 data set (Puck_190926_03). Vesalius was able to recover 28 uniform territories such as the embryonic liver and eye (Fig 2c). As expected, it is not easy to define clear territories with BayesSpace and Seurat as numerous small regions were identified (Fig. 2d, Fig. S5 and Fig. S6).

### Vesalius outperforms other Spatial domains tools in heterogenous territories

To assess the performance of Vesalius, we simulated ST data with 3 territories under three different regimes (See Methods – Fig. S1c): *pure, uniform* and *exponential*.

The *pure* regime contains a single cell type in each territory. The *uniform* regime contains *n* different cell types in each territory and each cell type appears in equal proportion. Finally, the *exponential* regime contains *n* cell types and the overall proportion of each cell type changes between territories following an exponential pattern. For uniform and exponential regime, we evaluated the performance of each tool with n = 3,4, and 5 cell types in each territory. Simulated data sets contain ∼6000 cells divided between 3 equally sized territories.

In these simulations, Vesalius generally recovered the 3 territories successfully across all conditions (Fig 2e – Fig. S7 -S16). Vesalius can clearly distinguish territories containing single cell types (pure), different cell types (uniform) or differences in cell type proportions (exponential). BayesSpace only recovered three clear territories in the pure regime and not in all simulation replicates (Fig 2e – Fig. S7 -S16). In the uniform and exponential regimes, BayesSpace often misattributed barcodes or in the more extreme cases only recovered two territories instead of three. As expected, Seurat did not recover territories but rather cell type patches (Fig 2e – Fig. S7 -S16). Interestingly, both BayesSpace and Seurat displayed gradual changes cluster locations in the exponential regime. However, only Vesalius can clearly differentiate territories with varying cell type proportions. It is remarkable that Giotto did not recover any territory even in the pure regime and only identified numerous small patches.

To quantitively assess the performance of Vesalius compared to other tools, we used an Adjusted Rand Index (ARI)^23^. While ARI scores are often used to compare clustering performance, they may be affected by cluster granularity and as such we also compared the performance of each tool using a Variation of Information metric^24^. Overall, Vesalius’s performance far exceeds its competitors (Fig. 2f) and it is also remarkable that the execution time for Vesalius is more than 10 times faster than BayesSpace and similar to the running time for Seurat (Fig. 2g).

### Vesalius illustrates tissue territories in sub-cellular resolution ST data

The challenge of heterogeneity is further amplified in sub-cellular resolution ST data as the transcriptome of each cell will likely be spread between multiple barcodes. Seq-Scope^8^ is an ST assay with a resolution smaller than 1um. To test if we can still recover territories, we applied Vesalius to murine Seq-Scope^8^ data in early-onset liver failure and crypt-surface colon (Fig. 2h-i). Vesalius illustrates hepatocellular zonation with both pericentral hepatocytes (Hep. PC) and periportal hepatocytes (Hep. PP) (Fig. 2h). We were also able to recover a territory of injured hepatocytes (Injured Hep.). Deciphering of tissue structure proved to also be successful in the colon where Vesalius shows territories related to smooth muscle, the crypt base and the crypt surface (Fig. 2i). Interestingly, our results suggest the emergence of different layers within the smooth muscle and crypt base.

### Isolating territories highlights the finer details of spatial patterning

We have shown that Vesalius is able to recover and isolate tissue territories from high resolution ST data especially tissue territories containing heterogenous cell populations. The isolation of territories enables an in-depth view of spatial patterning uncovering subtle expression patterns, new anatomical compartments and spatially resolved transcriptional shifts.

For instance, the analysis of the isolated CA field recovers all three CA field cell types namely CA1 pyramidal cells, CA2 pyramidal cells and CA3 pyramidal cells (Fig. 3a). In contrast, BayesSpace and Seurat were unable to recover all three sections (Fig 2b). We identified *Pcp4, Rgs14* and *Necab2* as marker genes for CA2, which is consistent with recent proteomics study for CA2 against CA1^26^. The *in situ* hybridization (ISH) images against these genes taken from the Allen Brain Atlas^22^ validate our prediction (Fig. 3b, Fig. S17). Further investigation of *Pcp4* expression illustrates that this gene has a strong expression in the thalamus and a comparatively weak expression in the CA2 field (Fig. 3c) and territory isolation by Vesalius enabled detection of the CA2 field in this noisy data.

**Fig. 3.**
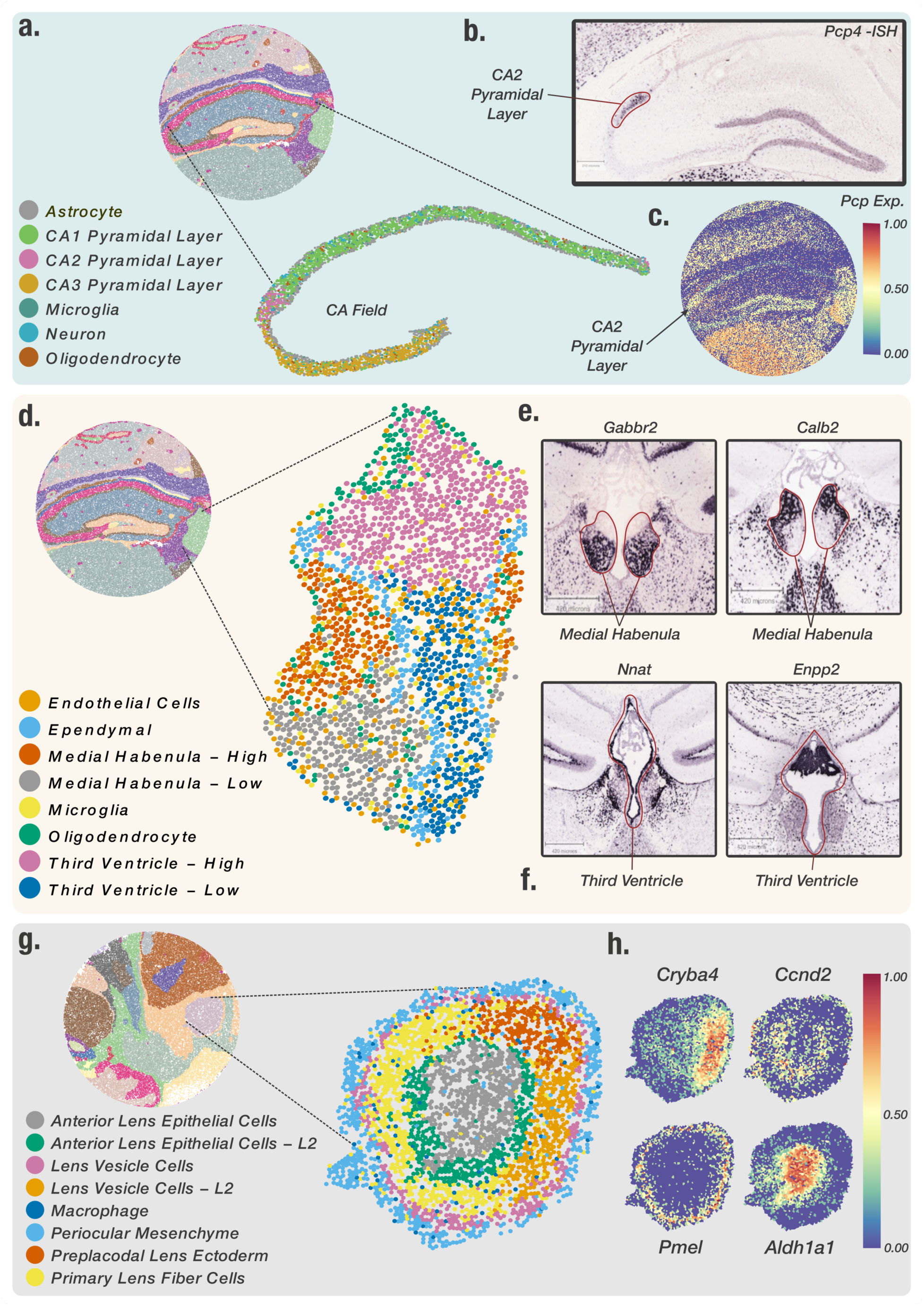
In depth analysis of isolated territories reveals the finer details of spatial patterning. (**a**) Mapping of clustered barcodes in the isolated CA field. Vesalius recovers all 3 CA pyramidal layers. (**b**) CA2 pyramidal layer was enriched with, among others, *Pcp4* a canonical CA2 layer marker. The ISH image taken from the Allen Brain Atlas corroborates the positioning of the CA2 layer within the isolated CA field. (**c**) *Pcp4* expression within the CA2 layer is lost in favor of stronger expression in the thalamus. (**d**) The isolated medial habenula and third ventricle show distinct spatial compartments after barcode clustering. (**e**) ISH image describing a medial habenula lower compartment marker (*Gabbr2*), and a medial habenula upper compartment marker (*Calb2*). The medial habenula is highlighted in red. (**f**) ISH images of lower third ventricle marker (*Nnat*) and upper third ventricle marker (*Enpp2*). (**g**) Barcode clustering and mapping of the embryonic eye (E12.5) show that Anterior Lens Epithelial cells and Lens Vesicle cells are separated into distinct layers. (**j**) Differential gene expression analysis between Anterior Lens Epithelial Cell layers revealed that the expression of *Cryba4* was restricted to the inner layer while *Cnnd2* was expressed in the outer layer. Similarly, *Pmel* was expressed in the outer Lens Vesicle cell layer and *Aldh1a1* was expressed in the inner layer.

The isolation of the third ventricle and the medial habenula further demonstrates how subtle gene expression patterns can be retrieved by using Vesalius. Indeed, the medial habenula and the third ventricle are divided into distinct spatial compartments (Fig. 3d). We observed an upper and lower medial habenula compartment characterized by distinct gene expression patterns. Overall, 119 genes were differentially expressed (p<0.05) between both compartments (Supplementary Table 3-4). For example, *Gabbr2* showed a higher expression in the lower medial habenula compartment (Fig. 3e – left) while *Calb2* showed a higher expression in the upper compartment (Fig. 3e - right). The medial habenula is highlighted in red. Similarly, the third ventricle exhibited 288 differentially expressed (p<0.05) between the upper compartment and lower compartment (Supplementary Table 3-4). *Nnat* is more strongly expressed in the lower third ventricle (Fig. 3f - left) and its expression pattern coincides with the ependymal cell layer that lines the ventricular system. By contrast, the expression of *Enpp2* was located in the upper third ventricle (Fig. 3f - right) and is absent from the ependymal cell layer. The third ventricle is highlighted in red. Interestingly, we also found a distinct cluster of ependymal cells lining the third ventricle (Fig. 3d).

In Slide-seq V2 embryo data, the developing eye displayed subtle transcriptional patterning (Fig. 3g). Anterior Lens Epithelial Cells are characterized by two distinct spatial patterns organized in a concentric fashion. We observed a similar effect in Lens Vesicle cell distribution. We compared gene expression between Anterior Lens Epithelial cell layers and found 81 differentially expressed genes (Table S2). For example, *Cryba4* was highly expressed in the inner layer while *Ccnd2* was expressed in the outer layer (Fig. 3h – top row). Similarly, differential gene expression analysis between Lens Vesicle cells returned 74 differentially expressed genes including *Pmel* and *Aldh1a1* (Fig. 3h).

While it still remains unclear if these shifts represent novel cell types in the developing eye or spatially resolved developmental cues, Vesalius provides an easy, robust and reproducible way of accessing this information via territory isolation in high resolution ST.

### Cells show territory specific gene expression in the mouse hippocampus

Territory isolation enables us to study territory-specific genes for the same cell type. To ensure that we are only accounting for the transcriptome of a single cell type, we first ran Robust Cell Type Decomposition^27^ (RCTD) on Slide-seq V2 mouse hippocampus data (See Methods). We selected all barcodes (n=12 013) that contained a single cell type or homotypic spots and used these annotations to assign cell types across the mouse hippocampus. Next, we isolated territories corresponding to the cortex (1 to 6 and 8 in Fig. 2a) and thalamus (28 and 33 in Fig. 2a). We used Vesalius to compare cells that appear in both territories (> 30 cells) and discovered 73 differentially expressed genes (p <0.01) across 4 cell types (Astrocytes, Interneuron, Oligodendrocytes, and Entorhinal cells) (Supplementary Table 4). We illustrate the differential expression of *Cpe* in Astrocytes and *Nrgn* in Entorhinal cells in Figure 4a. Astrocytes exhibit a stronger expression of *Cpe* in the thalamus compared to the cortex while Entorhinal cells show a higher expression of *Nrgn* in the cortex.

**Fig. 4.**
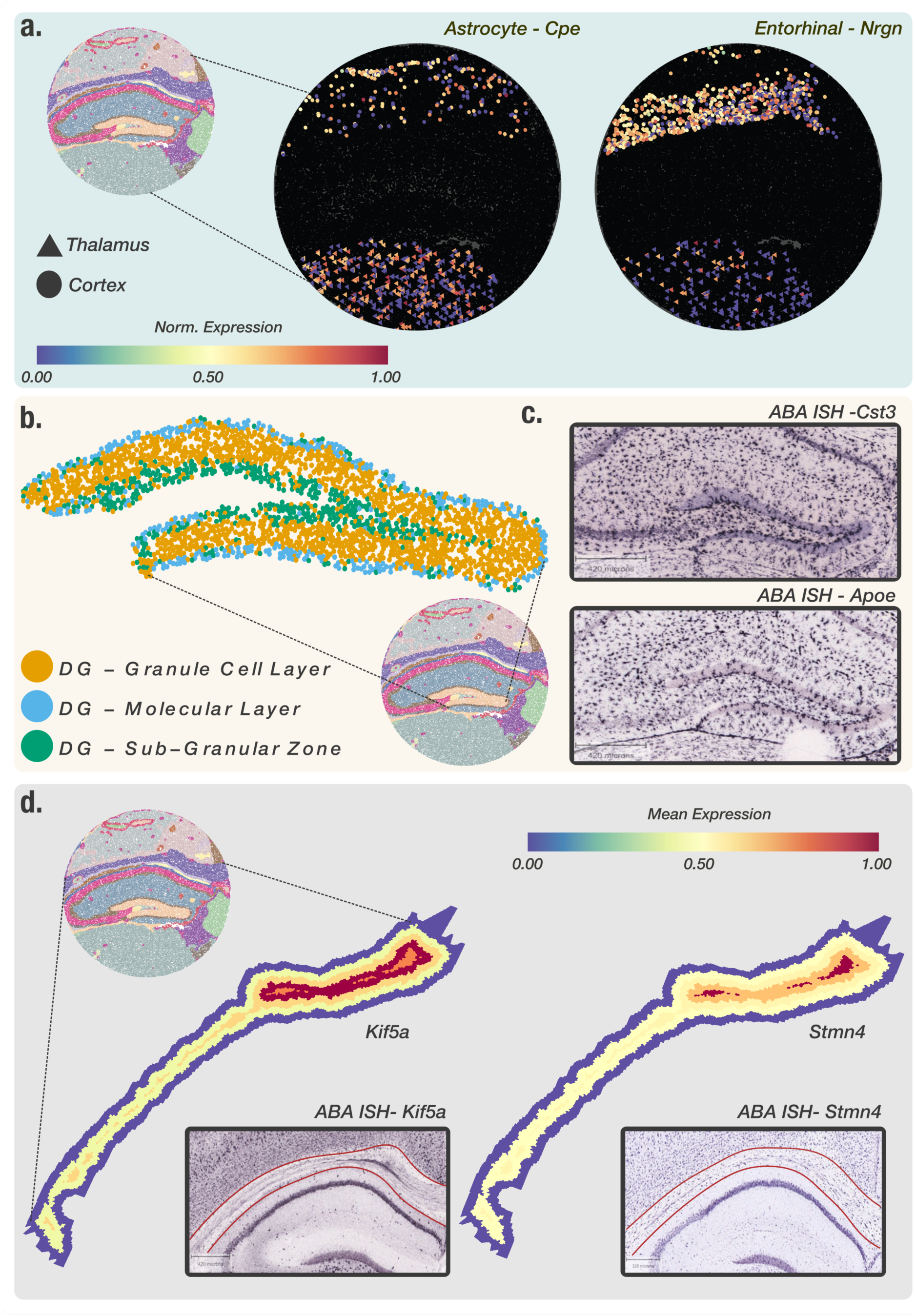
Investigation of territories reveals gene expression patterns linked to neighboring tissues and tissue morphology. (**a**) Differential gene expression analysis between cells contained in the cortex and the thalamus shows that spatial location influences gene expression. Astrocytes in the cortex are enriched with *Cpe* while Entorhinal cells in the thalamus are enriched with *Nrgn*.(**b**) Barcode clustering of the isolated dentate gyrus reveals transcriptional dissimilarity between each Dentate Gyrus (DG) layer. (**C**) Differential gene expression analysis between DG Granule cell layer and DG sub granular zone displayed a high expression of *Cst3* and *Apoe* at the border between layers. *Cst3* and *Apoe* border expression is corroborated by Allen brain Atlas ISH images. (**D**) Layered expression pattern of *Stmn4* and *Kif5a* within the isolated Corpus Callosum showed a higher expression at the center of the Corpus callosum. ISH images corroborate the spatial expression pattern of both genes. Corpus callosum contained within red lines.

### Investigation of territories reveals gene expression patterns linked to neighboring tissues and tissue morphology

We also tested if gene expression patterns could be linked to the morphology of a territory. Each territory provided by Vesalius can be manipulated using morphological operators (See Methods) or even divided into layers. The use of morphological operators for example permits the inclusion of additional barcodes that are part of the neighboring tissue. A territory divided into layers can be used to compare gene expression between the center of the structure and the edge of the structure.

Inflation and barcode clustering of the Dentate Gyrus (DG-GCL) revealed that this territory included a thin layer of barcodes belonging to the Dentate Gyrus – Sub-granular zone (DG-SGZ) and the Dentate Gyrus – Molecular layer (DG -ML) (Fig. 4b). Differential gene expression analysis displayed two genes *Cst3* and *Apoe* expressed at the border between the DG-GCL and the DG-SGZ (See Table S3). Allen Brain Atlas ISH images corroborate these results by displaying a higher expression of both genes at the border of the DG-GCL and the DG-SGZ (Fig. 4c). While increased expression of *Cst3* and *Apoe* could results from a high cellular density at the border, our results demonstrate how Vesalius can aid in recovering subtle spatially driven gene expression or cellular patterns at the border between tissues.

We also isolated, dilated and performed a layer analysis of the Corpus callosum. Remarkably, we found high expression of *Stmn4* and *Kif5a* in the inner most layers (Fig. 4d). Mean expression of both genes gradually increased towards the core of the corpus callosum. The ISH images for *Stmn4* taken from the Allen Brain Atlas showed a stripe of *Stmn4* expression (Fig. 4d). *Kif5a* also shows a stripe and strong expression towards the center of the murine brain. The corpus callosum is highlighted in red. All differentially expressed genes are listed in the Supplementary materials.

## Discussion

Spatial transcriptomics - as a technique - has demonstrated its ability to recover the transcriptome of cells within the context of a tissue^2^. This technical advancement has highlighted the influence of spatial context and cellular micro-environment on the transcriptome of cells^29^. Alongside experimental development, computational frameworks have been developed in order to analyze these new data sets^11–17^. However, it still remains challenging to accurately recover spatial domains especially when these domains contain a multitude of cell types. Here, we present Vesalius – an R package to effectively perform *in silico* anatomization of high-resolution ST data. Vesalius provides a reproducible framework for the isolation and in-depth analysis of tissue territories.

The key to Vesalius’s effectiveness in recovering heterogenous tissue territories resides in the embedding of the transcriptome into RGB colored images (Fig 1a – Fig S1a). The application of image processing techniques to these images retrieves tissue territories without any assumption about the transcriptome of neighboring cells. In this context, an anatomical structure is characterized by its transcriptome and overall cell composition. Taken together, Vesalius can easily segment tissue territories containing heterogenous cell populations (Fig 2a-c).

In contrast, computational tools such as BayesSpace^12^ and Giotto^13^ assume that neighboring spots are likely to be part of the same cluster if they are transcriptionally similar. While this assumption holds in lower resolution or in homogenous tissues, high resolution ST data sets recover the cellular complexity of tissues and thus neighboring spots are not always guaranteed to be of the same cell type. For example, BayesSpace is able to recover anatomical structures in Slide-Seq V2 as long as these structures are composed of similar cell types (Fig. 2b-d – Fig. S3-6). These results are comparable to those produced with Seurat (Fig 2b-d - Fig. S3-6) that does not consider the spatial component at all and simply maps cell cluster to their respective locations. The current trend in sequencing-based ST to increase resolution^8,30–32^ and ultimately achieving sub-cellular resolution will prove extremely challenging to analyze if heterogeneity is not considered carefully.

We further demonstrate the challenge of dealing with heterogenous territories by benchmarking BayesSpace, Giotto, Seurat and Vesalius in simulated data sets. Our simulations were designed to include different scenarios that could arise from real high-resolution ST data (Fig. S1c). Overall, Vesalius is the only algorithm able to detect territories in all regimes (Fig 2e-g – Fig. S7-16). In all cases, Giotto was unable to recover any clear territory even in the pure regime. Seurat also falters but this reflects the fact that Seurat was never designed to recover spatial domains accurately but rather cluster similar transcriptomes together. BayesSpace did not identify clear separations between territories once heterogeneity was introduced. Vesalius can recover territories much more accurately, especially in cases where a territory contains multiple cell types or each territory is characterized by a shift in cell type proportion.

Isolation of spatial territories enhances the analysis of ST data. The clusters provided by BayesSpace and Seurat do not provide a convenient way of comparing spatial territories (Fig. S3-S6) and sub-clustering does not equate to spatial domain analysis due to the wide spread of barcodes within clusters. We exemplify how the isolation of uniform territories provides an easy and reproducible way of investigating the finer details of spatial patterning. The CA2 field is often lost and merged with the CA1 and CA3 field (Fig. 2b) yet we were able to recover all three CA fields in the mouse hippocampus after territory isolation (Fig. 3a). The isolation of territories ensures that weaker expression patterns such as the expression of *Pcp4* – a CA2 field marker - are not lost in favor of stronger ones (Fig 3b-c). This approach also illustrated that the medial habenula and third ventricle were in fact divided into compartments (Fig 3d). For example, we showed that 119 genes are differentially expressed between medial habenula compartments with the expression of *Gabbr2* and *Calb2* delineating the position of both compartments in ISH images taken from the Allen Brain Atlas (Fig 3e). The isolation of the embryonic eye (Fig. 3g-h) displayed transcriptional shifts that could either indicate novel cell types or spatially resolved developmental cues occurring during development. Finally, we were able to use isolated territories and demonstrate that cells in the mouse hippocampus exhibit territory specific gene expression patterns: Astrocytes in the cortex showed a higher expression of *Cpe* compared to astrocytes in the thalamus (Fig. 4a). These results demonstrate how crucial spatial context is to understand the transcriptome of cells.

Vesalius’s unique way of representing territories also lends itself to the manipulation of territories and investigating gene expression in the context of neighboring tissues. For instance, we identified an increased expression of *Cst3* and *Apoe* at border of the DG-GCL and DG-SGZ. We also discovered a heightened expression of *Kif5a* and *Stmn4* at the center of the corpus callosum (Fig. 4d).

Overall, Vesalius is an effective tool to recover spatial domains in high resolution ST data and isolate territories for further analysis. We demonstrate that the isolation of spatial domains uncovers the finer details of spatial patterning and capitalizes on the wealth of information contained in high resolution ST data. Vesalius complements other ST analysis methods and enhances potential biological insights.

## Materials and Methods

### Pre-processing

Pre-processing of data prior to Vesalius image building was handled by the Seurat package^11^. Slide-seq V2 data sets were loaded, log normalized and scaled using the default Seurat settings. Variable features (n=2000) were extracted prior to PCA. We used log normalization as the goal was to acquire general tissue territories for further analysis. Seq-Scope data sets come already pre-processed.

### Cell type deconvolution

To infer the cell types of individual beads in the Slide-seq V2 data sets, RCTD (Robust Cell Type Decomposition) was used^27^. RCTD is a R package that decomposes the cell type mixture of spatial beads with resolution lower than a single cell. It is trained with single cell RNA-sequencing (scRNA-seq) data sets with annotated cell types. Based this, RCTD predicts the cell types of each bead. As RCTD was specialized to spatial transcriptomics data with fine resolution such as Slide-seq, we selected it as a deconvolution tool. We conducted simulation under doublet mode, which is recommended for Slide-seq data sets^27^. It assumes that each bead contains up to two different cell types. We trained RCTD using scRNA-seq data set for hippocampus^33^ that consists of 27,953 genes and 113,507 cells annotated with 17 cell types. The scRNA-seq data set was obtained from the publicly available repository, DropViz. Afterwards, we ran RCTD to infer the cell types of the beads in the Slide-seq V2 data sets. The results of the cell-type decomposition were validated by cell type markers accompanied with the scRNA-seq data set from hippocampus^28^.

### Vesalius – Transcriptome embedding into the RGB color space

To embed the transcriptome into the RGB color space, Vesalius first computes Principal component analysis (PCA) followed by a 3-dimensional Uniform Manifold Approximation and Projection (UMAP). UMAP latent space is min-max normalized for each barcode *Bi:*

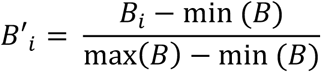

The normalized 3-dimesional space can simply be converted into an RGB color space (Fig. S1a). Alternatively, Vesalius can directly use PCA loading values in which case Vesalius selects a PC slice composed of 3 principal components – one for each color channel (RGB). For each color channel *c* and for each barcode *B*_*i*_, Vesalius takes the sum of the absolute value of all loading values associated to *B*_*i*_ in color channel *c*:

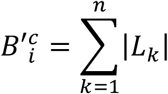

With *k= 1…n* and *n* being the number of non-zero loading values associated with barcode 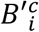.

For each color channel, Vesalius returns a numerical array of length equal to the number of barcodes. To ensure that these values are within color space bounds, each color channel is min-max normalized. As three PCs are selected in a “slice” (e.g. PC1, PC2, and PC3) and RGB color space contains three dimensions, Vesalius simply assigns the color value obtained from each PC into each color space dimension (Red, Green, and Blue – Fig. S1a).

### Vesalius – Building images arrays from Spatial transcriptomics

Each ST data set contains a set of coordinates describing the x and y position of each barcode/spot. As the coordinates will not necessarily be uniformly distributed and contiguous, Vesalius expands punctual coordinate values into tiles using Voronoi tessellation and fills each tile using rasterization. First, we filter stray barcodes by comparing barcode density in a grid covering the entire ST assay and removing any barcodes that fall into a grid section with a low barcode density (Determined based on quantiles of grid density). Next, we produce a Voronoi diagram using the remaining barcodes. Each tile is then converted into a pixel set via rasterization. The color code obtained for each barcode is then assigned to its tile. RGB image arrays are three dimensional arrays and as such each color channel in the image will receive a distinct set of color value for each latent space dimension (UMAP or PCA). The tiles associated to each barcode remain the same between color channels. Optionally, Vesalius can resize the image output using Nearest-neighbor interpolation as default. All barcodes will be retained after image resizing.

### Vesalius – Image processing and segmentation

Image arrays are handled by the *imager*^34^ and *imagerExtra* R packages which contain a set of image processing methods such as blurs, segmentation and image manipulation. Image can be regularized with Nonlinear total variation-based noise removal algorithms^35^. Vesalius provides an iterative segmentation approach to reduce the color space of an image and extract territories.

First, the image array is smoothed using either Gaussian blur, median blur, box blur or a combination of the aforementioned methods. Second, color values are clustered in *k* clusters using K-means clustering. The choice of *k* is left to the user and will depend on the level of territory refinement the user wishes to have. Multiple rounds of smoothing and segmentation may be applied. In the instance that multiple values of *k* are used, Vesalius will smooth and segment the image as describe above and repeat the process for all values of *k*. A decreasing *k* value at each round will iteratively decrease the color space.

To decrease computational time, the clustering is only applied to the center pixel value. The center pixel is defined by the pixel corresponding to the original barcode location before tessellation and rasterization. Vesalius also provides the option to smooth and segment images using all pixels instead. Using all pixels for segmentation produces sharper segments between homogenous territories. However, this sharpness in homogenous territories may come at the cost of increased noise in heterogenous territories.

### Vesalius – Isolating Territories

Once barcodes have been assigned to a color cluster, each color cluster is further subdivided into territories. For every barcode present in a color cluster, Vesalius checks for all barcodes that are within a certain capture radius of each other and assigns them to a territory (Fig. S1b). This process is repeated for all barcodes in a color cluster until they have all been assigned to a territory. Vesalius repeats this process for each color cluster. The capture radius is defined as the distance value that corresponds to the proportion of maximum distances between all barcodes in the ST assay. First, Vesalius computes a distance matrix between all barcodes in 2D space and extracts the maximum possible distance between barcodes. A user-selected parameter defines the proportion of this maximum distance that should be used as a capture radius.

While the number of color clusters is fixed by *k* (or final *k*) as described above, the final number of territories may vary depending on parameter selection used throughout the analysis. This includes images processing and territory isolation. The selection of these parameters depends on the users interests and how they wish to explore ST data.

### Simulating high resolution ST data and tissue heterogeneity

To benchmark Vesalius in high resolution ST data sets containing heterogenous tissues, we simulated ST data using Slide-seq V2 after cell type deconvolution. First, we applied RCTD^27^ to deconvolute cell types and selected barcodes that contained a single cell type. This approach ensures that we more closely mimic the biology of heterogenous tissue rather than ST assay technical limitations. Next, we generated 6000 random x and y coordinates pairs that can be divided into three equally sized territories. For each territory, we randomly sampled cell types and barcodes under three different regimes: pure, uniform and exponential (Fig. S1c). The pure regime contains a single randomly sampled cell type in each territory and randomly selected barcodes for that cell type. The uniform regime contains *n* cell types in equal proportion. Each territory contains different sets of randomly sampled cell types. Finally, the exponential regime contains *n* cell types in unequal proportion:

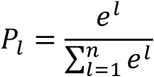

Where *n* is the number of different cell types and *P*_*l*_ is the proportion of cell type *l*. Each territory contains the same cell types in different proportions. To reduce any potential cell type bias, we ran the simulation 10 times each time randomly selecting different cell types and different barcodes. The simulated data set produced at each sampling round was the same between each tool and a summary of the cell types sampled at each round can be found in Supplementary Table 1. The code used to run simulations and method comparison (see below) can be found here: https://github.com/patrickCNMartin/Vesalius/tree/main/methodComp

### Method comparison

We compared the performance of Vesalius to the performance of other methods such as Seurat^11^, BayesSpace^12^ and Giotto^13^ in our simulated data sets.

BayesSpace required a minor modification to accommodate Slide-seq V2. We adapted the. *find_neighbors* function to select neighbors based on Euclidean distance in 2D space and select the 6 closest neighbors. We added a new platform named “SS” for Slide-Seq. The forked and modified version of BayesSpace is available at: https://github.com/patrickCNMartin/BayesSpace.

Simulated data contained 3 territories and as such each barcode was clearly assigned to a territory and provided us with a ground truth. We assessed each tool’s ability to recover each territory using and Adjusted Rand Index^23^ and Variation of information metric^24^. ARI scores are affected by the granularity of the clustering and the Variation of information metric provides an alternative measure of performance. It is important to acknowledge that despite the fact that Seurat is consistently used for spatial domain retrieval benchmarking, Seurat is not designed to retrieve spatial domains and is expected to perform poorly in highly heterogenous tissues.

All code related to the comparison between methods can be found here: https://github.com/patrickCNMartin/Vesalius/tree/main/methodComp

### Vesalius – Differential Gene expression and territory Markers

Marker genes and differentially expressed genes can be extracted from each territory. This process can be carried out in a fivefold manner:

- Territory VS all other territories combined
- Territory VS all other territories individually
- Territory(ies) VS territory(ies)
- Cell within a territory(ies) VS cell within a territory(ies)
- Layer in territory VS layer in territory

To be considered for differential expression analysis, genes must pass a set of criteria. First, genes must be present in a certain percentage of barcodes in at least one territory (>10% of beads as default). Second, the log fold change must be above a certain threshold (logFC >= 0.25). It should be noted that this threshold is applied in the case of up-regulation as well as down-regulation. Remaining genes are tested for significant differential gene expression by using a Wilcoxon rank-sum test (Bonferroni corrected p-value < 0.05). Gene expression patterns can be visualized by using the *viewGeneExpression* function provided in the Vesalius package. Gene expression can be visualized over the entire slide or in an isolated territory. For visualization, gene expression is min-max normalized.

### Vesalius – Territory dilation, erosion, filling and cleaning

By using image representation of territories, Vesalius provides a convenient way to manipulate territories using image morphology. Vesalius encompasses dilation, erosion, filling, and cleaning into a single function. We summarize image morphologies using a “morphology factor” described by an array of integers. Positive integers increase territory size while negative integers decrease territory size. Numerical arrays of positive and negative integers provide filling and cleaning morphologies. For example, in the case of a cleaning morphological operator, a morphology factor array *v* = [-5,5] will first erode the territory by 5 pixels and then dilate the territory by 5 pixels.

### Vesalius – Territory Layering and layered gene expression

Isolated territory layering is achieved by capturing territory edges and removing barcodes belonging to the edge and repeating the process until no more barcodes remain. First, the isolated territory is converted to a black and white image and X-Y Sobel edge detection is applied. All barcodes that share a pixel with the detected edge are pooled into a layer and removed from the territory. The edge detection and pooling process are repeated until all barcodes have been assigned to a layer. Layers can be combined by merging neighboring layers together (Fig. S1d). Differential gene expression between layers is carried out using a Wilcoxon rank-sum test (Bonferroni corrected p-value < 0.05 & logFC >= 0.25). Visualization of gene expression between layers is provided by the *viewLayeredExpression* in the Vesalius package. Layers are described by their normalized and averaged expression values.

### Cell Clustering and Annotation

Clustering analysis of isolated territories was carried out using the Seurat package. Clusters and territories were manually annotated using their respective genetic markers. Markers were extracted from each cluster using the *FindAllMarkers* function provided by Seurat and the *extractClusterMarkers* function provided by Vesalius. *FindAllMarkers* compares clusters between each other (in the isolated territory) while *extractClusterMarkers* compares clusters to all other barcodes present in the slide. This distinction ensures that we recover subtle differences between cell types as well as territory specific gene expression.

We used the default Wilcoxon Rank-sum test for marker extraction. Identified markers were compared to Allen Brain Atlas^22^ (https://mouse.brain-map.org/), the lifeMaps/geneCard database^36^ (https://discovery.lifemapsc.com/in-vivo-development) or panglaodb database^37^ (https://panglaodb.se/) to assign cell type to clusters and territories. Manual annotation of cell clusters was preferred over automated methods to ensure correct tissue annotations, rare cell type annotation and finally to maintain subtle spatially driven patterning. Cell type markers used for annotation are available in Supplementary Table 2.

## Acknowledgments

We would like to thank Konstantin Khodosevich for critical reading of the manuscript.

## Author contributions

Conceptualization: PCNM, KJW

Methodology: PCNM, BWH, KJW

Investigation: PCNM, HK, CL

Visualization: PCNM

Funding acquisition: KJW

Project administration: KJW

Supervision: KJW

Writing – original draft: PCNM, KJW

Writing – review & editing: PCNM, HK, CL, BWH, KJW

## Data availability

Slide-seq V2 data sets were downloaded from the Single Cell Portal : https://singlecell.broadinstitute.org/single_cell/study/SCP815/highly-sensitive-spatial-transcriptomics-at-near-cellular-resolution-with-slide-seqv2#study-summary

Seq-Scope data sets were downloaded from: https://deepblue.lib.umich.edu/data/concern/data_sets/9c67wn05f

All ISH images used for validation were taken from the Allen Brain Atlas: https://mouse.brain-map.org/

All differentially expressed genes and marker genes used in this analysis are available in the supplementary materials.

## Code availability

The Modified version of BayesSpace to accommodate Slide-seqV2 data is available here: https://github.com/patrickCNMartin/BayesSpace

Vesalius is available here: https://patrickcnmartin.github.io/Vesalius/index.html

The analysis presented in this manuscript is both described and discussed here: https://patrickcnmartin.github.io/Vesalius/articles/Vesalius_Analysis/Vesalius_analysis.html

The code related to the comparison and the simulation of high-resolution data is available here: https://github.com/patrickCNMartin/Vesalius/tree/main/methodComp

## Supplementary Figures

**Fig. S1.**
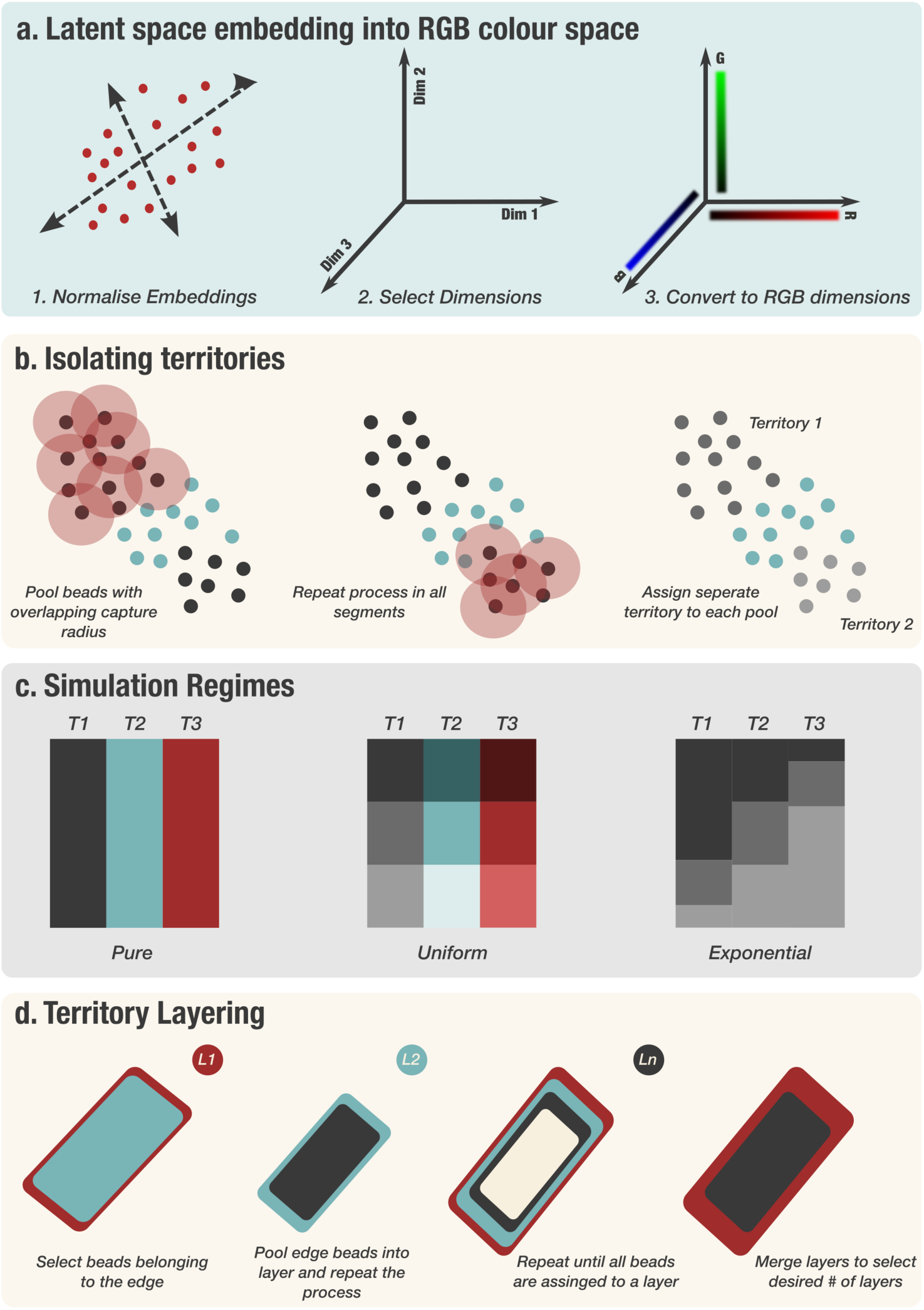
Vesalius methods visualized. (**a**) Vesalius converts the transcriptome of cells by using a normalized latent space (either UMAP or PCA) and using embedding values as RGB color values. (**b**) After image segmentation, Vesalius further isolates territories by pooling barcodes that are close to each other in 2D space. Vesalius finds all barcodes that are within a capture distance of each other (represented by red circle) and assigns a unique territory to all these barcodes. The same process is applied to all beads of that color segment until all beads have been pooled into a distinct spatial territory. (**c**) Simulation regimes used to benchmark Vesalius in high resolution ST data. The pure regime only contains one cell type per territory. The uniform regime contains *n* different cell types in equal proportion in each territory. The exponential regime contains *n* cell types in varying proportions between territories. The cell types are the same between territories. (**d**) Territory layering uses images representation of territory to iteratively select the edge of a territory and assign a layer value to that edge. Once all barcodes have been assigned to a layer, the number of layers can be reduced by merging neighboring layers.

**Fig. S2.**
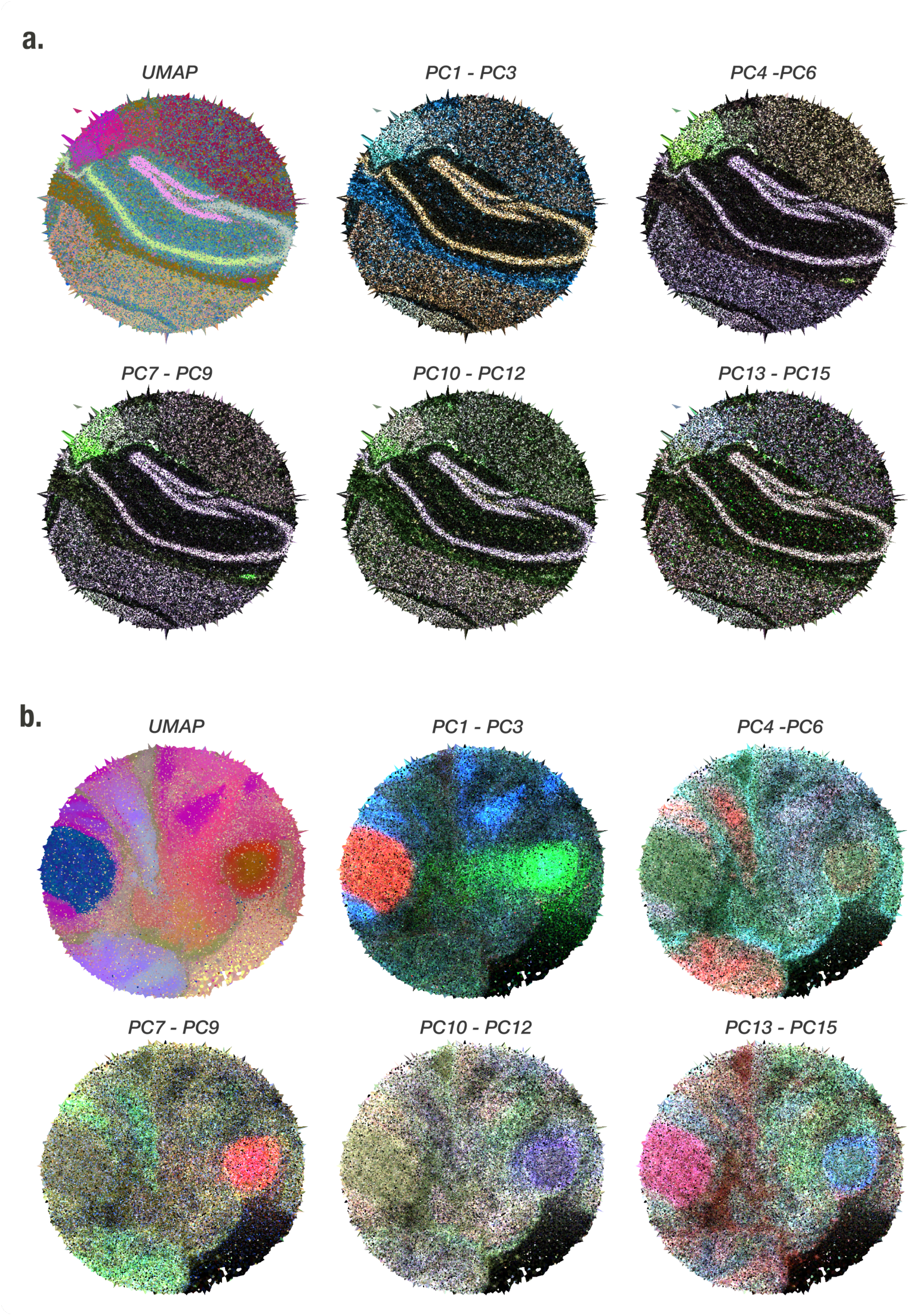
UMAP projection and PCA loading value define similar yet not identical territories. (**a**) UMAP and PCA slices in the mouse hippocampus (Puck_200115_08). Overall UMAP projections better recover the structure of the mouse hippocampus. (**b**) UMAP and PCA slices in the mouse embryo (Puck_190926_03). While UMAP projections recover more uniform territories, PCA slices enable a more targeted selection of territories. The selection of territories depends on the interest of the user and PCA enables more flexibility in the choice of territories.

**Fig. S3.**
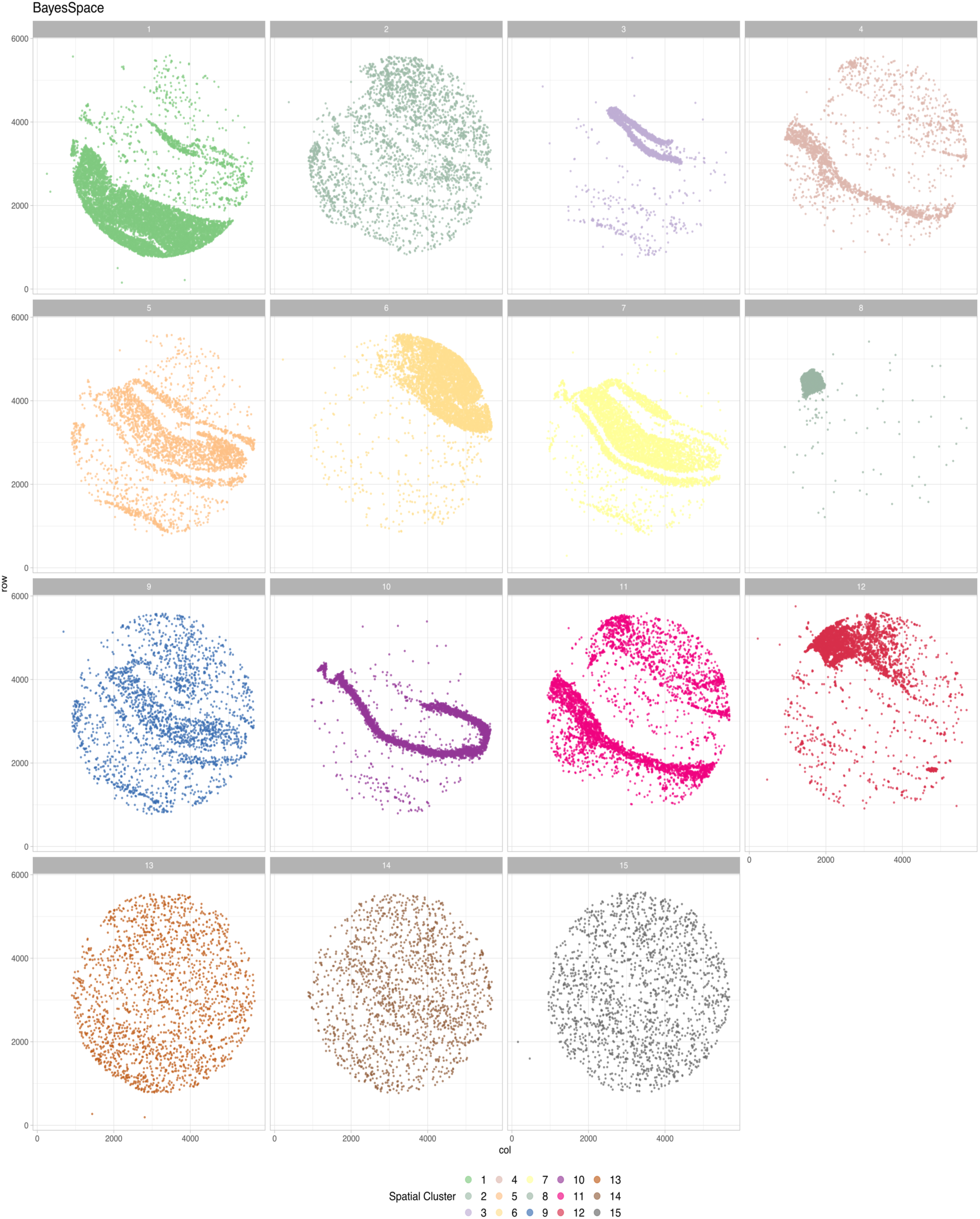
BayesSpace spatial cluster in mouse hippocampus. Visualizing BayesSpace clusters separately emphasizes that many of the spatial domain do not represent a specific anatomical structure. In this context, selecting a specific location in the tissue is impossible without either missing barcodes or by including unwanted barcodes. This also demonstrates why cluster sub-clustering does not equate to territory isolation and clustering.

**Fig. S4.**
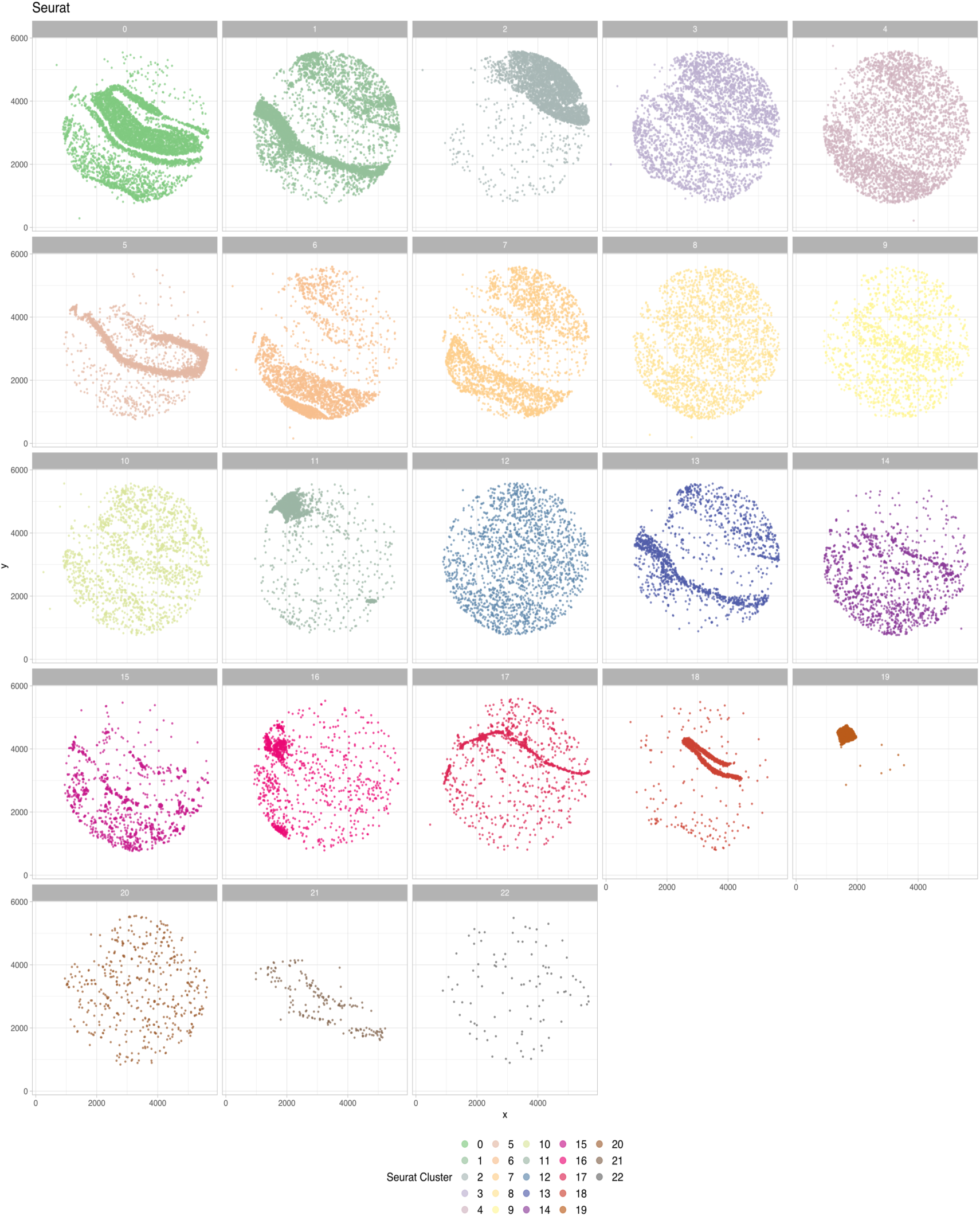
Seurat clusters in mouse hippocampus. Visualizing Seurat clusters separately emphasizes that many of the clusters do not represent a specific anatomical structure. In this context, selecting a specific location in the tissue is impossible without either missing barcodes or by including unwanted barcodes. This also demonstrates why cluster sub-clustering does not equate to territory isolation and clustering. Seurat is not designed to recover spatial domains but rather cluster transcriptional similarity.

**Fig. S5.**
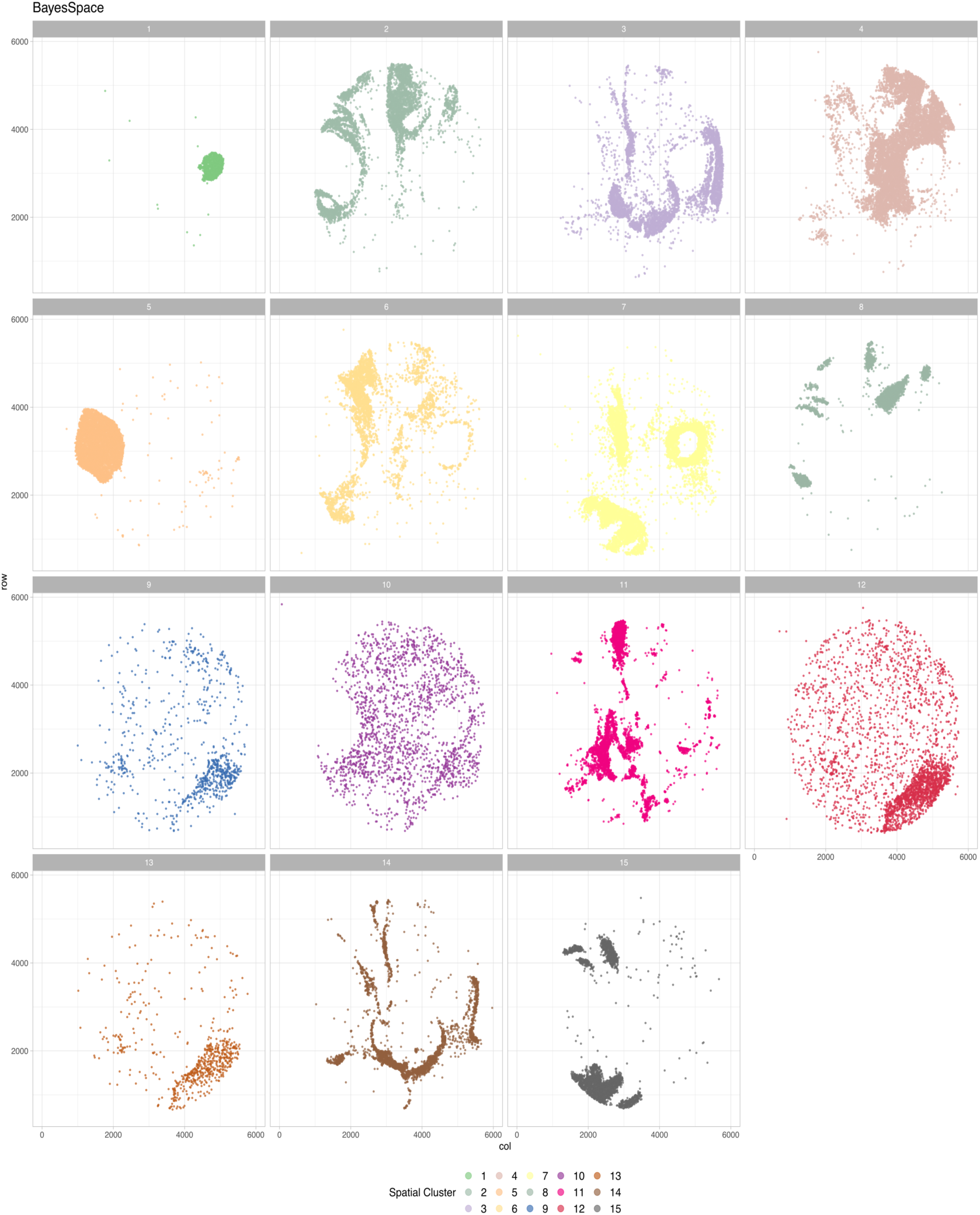
BayesSpace spatial cluster in mouse embryo. Visualizing BayesSpace clusters separately emphasizes that many of the spatial domain do not represent a specific anatomical structure. In this context, selecting a specific location in the tissue is impossible without either missing barcodes or by including unwanted barcodes. This also demonstrates why cluster sub-clustering does not equate to territory isolation and clustering.

**Fig. S6.**
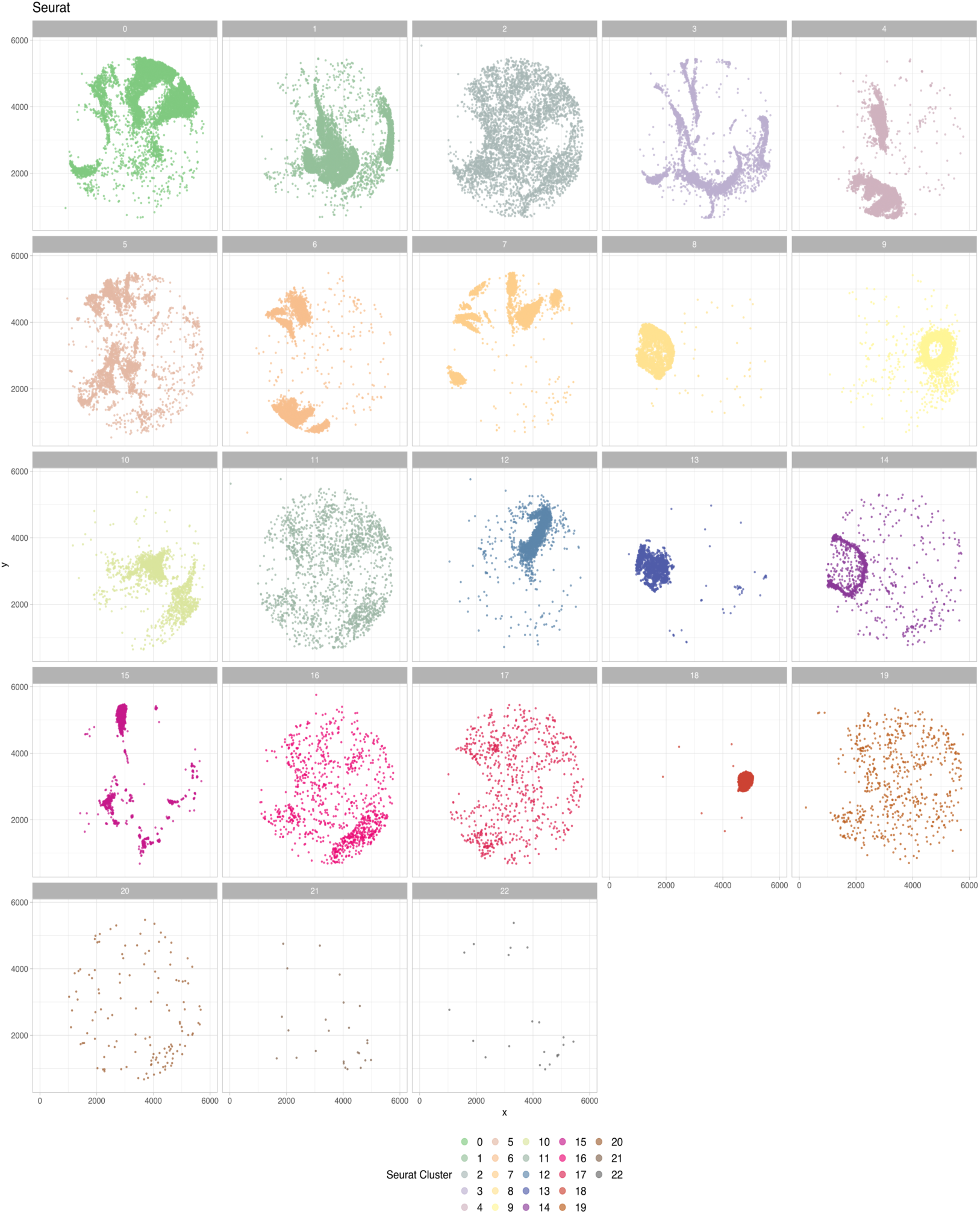
Seurat clusters in mouse embryo. Visualizing Seurat clusters separately emphasizes that many of the clusters do not represent a specific anatomical structure. In this context, selecting a specific location in the tissue is impossible without either missing barcodes or by including unwanted barcodes. This also demonstrates why cluster sub-clustering does not equate to territory isolation and clustering. Seurat is not designed to recover spatial domains but rather cluster transcriptional similarity.

**Fig. S17.**
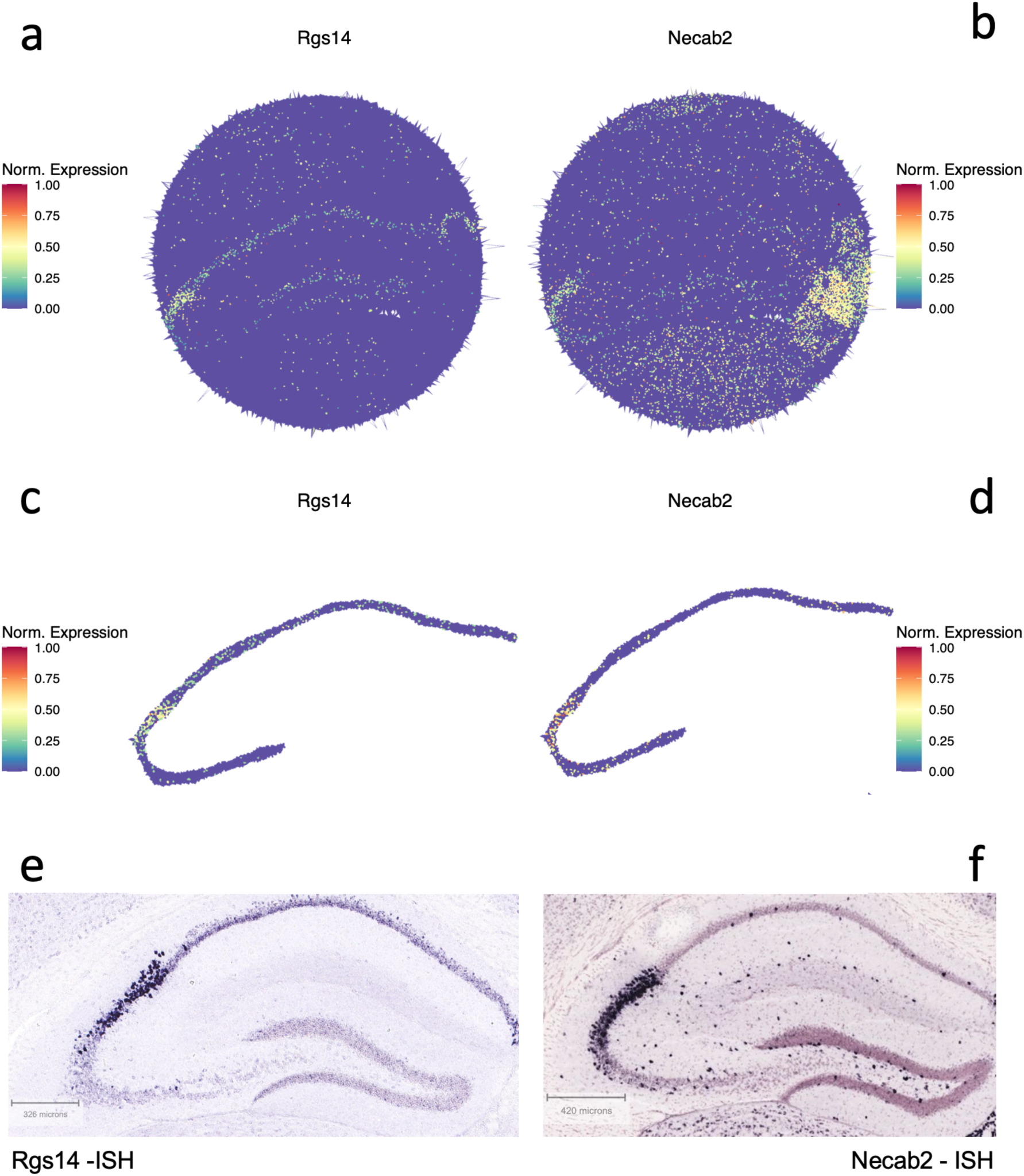
Validation of CA2 field Marker genes. Vesalius recovered the CA2 layer and canonical CA2 layer marker genes such as *Rgs14* and *Necab2*. (**A**) shows that the overall expression of *Rgs14* is fairly weak throughout the entire but increase in the CA2 layer. (**B**) *Necab2* is only weakly expressed in the CA2 field. (**C**) and (**D**) show how the isolated CA field emphasizes the expression of both marker genes. (**E**) and (**F**) confirm that both genes are indeed expressed in CA2 layer as described by the ISH images taken from the Allen Brain Atlas.

